# Regulation of reticular adhesions by KANK2 and talin2 in two melanoma cell lines

**DOI:** 10.1101/2025.06.02.657402

**Authors:** Anja Rac, Marija Lončarić, Nikolina Stojanović, Mahak Fatima, Mirna Rešetar, Dalibor Hršak, Jonathan D. Humphries, Martin J. Humphries, Andreja Ambriović-Ristov

## Abstract

Integrins bind to extracellular matrix proteins and, upon clustering, form multimolecular integrin adhesion complexes (IACs) that connect to and regulate the cell cytoskeleton, influencing various aspects of normal and tumour cell behaviour. Alongside well-characterized nascent adhesions, focal adhesions (FAs), fibrillar adhesions (FBs) and hemidesmosomes, a new class of IACs, reticular adhesions (RAs), have been identified. RAs, initially described as flat clathrin lattices formed by integrin αVβ5, lack association with actin and are devoid of FAs markers. The physiological role of RAs in normal and tumor cells is still incompletely understood and requires further investigation. Previously, we analysed IACs of two melanoma cell lines, MDA-MB-435S and RPMI-7951, grown under long term culture conditions, and demonstrated that both cell lines preferentially use integrin αVβ5 for adhesion. Here we present a comprehensive analysis of RAs in these two melanoma cell lines that differ in their ability to form FBs. To determine RAs composition, we treated cells with actin polymerisation inhibitor cytochalasin D (CytoD) which disrupts FAs, allowing isolation of RAs, which were analysed by MS-based proteomics, Western blotting and immunofluorescence. Known RA-associated proteins, including the AP-2 adaptor complex, disabled homolog 2 (DAB2) and Numb were identified in both lines, along with talin2. Notably, we also detected the presence of KN motif and ankyrin repeat domains protein (KANK2) in RA isolates. Proximity ligation analysis following CytoD-induced actin disruption confirmed the proximity of KANK2 and talin2 in RAs. We then investigated the effect of talin2 or KANK2 knockdown on RAs composition. While both talin2 and KANK2 are located in RAs, neither is essential for RA formation. Talin2 knockdown led to a reduction in RA components abundance in both cell lines. In MDA-MB-435S cell line, KANK2 produced a similar effect, mirroring the functional interaction of talin2 and KANK2 in FAs. However, in RPMI-7951 cells, KANK2 knockdown had no significant effect on RA components abundance. This discrepancy likely reflects the preferential localization of KANK2 in FBs and underscores the differing roles of talin2 and KANK2 in αVβ5-mediated FAs across the two cell lines. These findings underscore the complexity of adhesion signalling and highlight the importance of adhesion crosstalk in regulating cellular function.

## INTRODUCTION

Most cells in multicellular organisms require attachment to their surroundings through cell-cell or cell-extracellular matrix (ECM) adhesions. In cell-matrix adhesions, integrins, a family of transmembrane receptors, have long been recognized as the central scaffolds around which these complexes are built. Through binding to the ECM and clustering, integrins form multimolecular adhesion complexes (Chastney et al., 2021). The composition of integrin adhesion complexes (IACs) varies depending on the type of adhesions they form. Focal adhesions (FAs) mature from nascent adhesions and facilitate force transduction through tight links to contractile actomyosin fibers (Yamaguchi and Knaut, 2022, Mishra and Manavathi, 2021). Formed by many integrin heterodimers, FAs are located primarily at the cell edge. Fibrillar adhesions (FBs), formed by integrin α5β1 (Conway and Jacquemet, 2019), are located in the cell center of the migrating cell (Katz et al., 2000, Pankov et al., 2000). FBs arise through translocation of fibronectin-bound α5β1 integrins from the periphery to the center (Pankov et al., 2000, Clark et al., 2005) in a tension-dependent manner and have a role in fibronectin fibrillogenesis (Zamir et al., 2000, Lu et al., 2020). Integrins are also components of adhesion structures that lack strong actomyosin connections, are enriched in endocytic proteins (Lock et al., 2018, Zuidema et al., 2018), yet still function as signalling platforms (Leyton-Puig et al., 2017, Alfonzo-Mendez et al., 2022). These structures, formed as a result of frustrated endocytosis (Baschieri et al., 2018, Bruna-Gauchoux and Montagnac, 2022, Baschieri et al., 2020), and are referred to as reticular adhesions (RAs) (Lock et al., 2018), flat clathrin lattices (Baschieri et al., 2018, De Deyne et al., 1998, Zuidema et al., 2018, Leyton-Puig et al., 2017), or clathrin plaques (Moulay et al., 2020, Bucher et al., 2018, Lampe et al., 2016, Saffarian et al., 2009). RAs are mediated exclusively by integrin αVβ5 but lack most IAC components, except talin2 and tensin3 (Lock et al., 2018). Recent studies have shown that RAs are larger than flat clathrin lattices and contain them in an interspersed manner (Hakanpaa et al., 2023). Unlike FAs, RAs persist throughout cell division to enable effective mitosis and also transmit spatial memory from pre-mitotic to post-mitotic daughter cells (Zuidema et al., 2022, Lock et al., 2018). Since RAs form through arrested endocytosis, their core components include proteins involved in clathrin-mediated endocytosis, such as adaptor protein 2 (AP2), disabled protein 2 (Dab2), NUMB Endocytic Adaptor Protein (NUMB), Epidermal Growth Factor Receptor Pathway Substrate 15 (EPS15), Epidermal Growth Factor Receptor Pathway Substrate 15 Like 1 (EPS15L1), intersectin (ITSN), and Low Density Lipoprotein Receptor Adaptor Protein 1 (LDLRAP1 or ARH) (Lock et al., 2018, Baschieri et al., 2018, Zuidema et al., 2018, Lock et al., 2019). None of these proteins are present exclusively in RAs, and therefore cannot be used as markers. Lukas et al. (Lukas et al., 2024) have recently identified stonin1 as a protein exclusively present at RAs. However, they emphasized that stonin1 is not ubiquitously expressed.

Talins are key cytoplasmic mechanosensitive proteins that mediate binding of integrins to the cytoskeleton (Klapholz and Brown, 2017). There are two different isoforms of talin: talin1 and talin2. Talin1 forms the core of FAs, increasing the affinity of integrin for ligands (integrin activation) and recruiting a range of proteins (del Rio et al., 2009, Goult et al., 2021). Abrogation of talin1 expression leads to FA disruption, while talin2 has been shown to influence FA dynamics (Loncaric et al., 2023, Stojanović et al., 2025). Talin2 is mainly found at large FAs and FBs (Praekelt et al., 2012, Atherton et al., 2022) but also in RAs (Lock et al., 2018). Talins directly bind to actin and coordinate microtubule (MT) recruitment to adhesion sites via interaction with KANK proteins (Bouchet et al., 2016, Goult et al., 2018, Klapholz and Brown, 2017, Sun et al., 2016, Loncaric et al., 2023).

KANK2, a member of the KANK (kidney ankyrin repeat-containing) family proteins that bind the talin rod through a KN motif, is mainly found in mesenchymal cells (Chen et al., 2018). Along with KANK1, KANK2 localizes to the rims of mature integrin-containing FAs and adjacent regions called cortical microtubule stabilizing complexes (CMSCs). The CMSC, recruited via KANK2-talin1 interaction (Sun et al., 2016) or talin2-KANK2 interaction (Loncaric et al., 2023) stabilizes MTs near mature FAs and regulates actin-microtubule crosstalk. In some cells, behind the lamella, KANK2 binding to talin promotes integrin activation, and at the same time diminishes F-actin binding to talin. Consequently, this leads to the translocation of β1 integrins into FBs (Sun et al., 2016). FBs contain talin 2 (Praekelt et al., 2012) and KANK2 (Atherton et al., 2022, Stojanović et al., 2025).

RAs have been observed in various cell lines (Lukas et al., 2024, Lock et al., 2018, Paradzik et al., 2020, Zuidema et al., 2018); however, information on their molecular composition has mostly come from U2OS cells (Lock et al., 2018), keratinocytes (Zuidema et al., 2018), and C2C12 cells (Lukas et al., 2024). Here, a comprehensive analysis of RAs in two melanoma cell lines MDA-MB-435S and RPMI-7951, which differ in their ability to form FBs in long-term culture, is presented. In RAs of both cell lines, previously known RA proteins, including talin2 (Lock et al., 2018), were identified, as was KANK2. Although these two proteins were located in RAs, they were not essential for RA formation. Talin2 knockdown reduced the abundance of RA components in both cell lines. In the MDA-MB-435S cell line, KANK2 which functionally interacts with talin2 in FAs, produced a similar effect. However, KANK2 knockdown in RPMI-7951 cells did not affect the abundance of RA components. This difference between the MDA-MB-435S and RPMI-7951 cell lines is likely due to the preferential localization of KANK2 in FBs.

## MATERIALS AND METHODS

### Cell Cultures

The human melanoma cell lines MDA-MB-435S and RPMI-7951 were obtained from the American Type Culture Collection (ATCC). Cells were grown on uncoated surfaces in DMEM (Invitrogen) supplemented with 10% (v/v) FBS (Invitrogen) (DMEM-FBS) at 37°C with 5% CO2 (v/v) in a humidified atmosphere.

### Transient siRNA Transfection

For transient siRNA transfection experiments, cells (MDA-MB-435S, 4 × 10^4^, 4 × 10^5^ or 1.7 × 10^6^; RPMI-7951, 3.7 × 10^4^, 5 × 10^5^ or 1 × 10^6^) were plated in 24-well, 6-well cell culture plates or 10 cm diameter Petri dishes to achieve 60–80% confluence after 24 hours. Cells were transfected 24 hours later, using Lipofectamine RNAiMax (13778150, Thermo Fisher Scientific), with 25 nM of control (Silencer™ Select Negative Control No. 1 siRNA, Ambion), ITGA5 (s7547, Ambion), ITGB5 (s7591, Ambion) or gene-specific siRNA for KANK2 (target sequence: ATGTCAACGTGCAAGATGA), TLN2 (target sequence: TTTCGTTTTCATCTACTCCTT) (Loncaric et al., 2023), all purchased from Sigma. Knockdown was validated by SDS-PAGE and western blotting (WB) using specific antibodies and matched labelled secondary antibodies (Additional file 1 Table S1).

### Cell treatment

To enrich samples with RA proteins, cells were treated with different concentrations (1, 4, 7, 20 nM in DMSO) of actin polymerization inhibitor cytochalasin D (CytoD) for 2 hours at 37°C. Control cells were treated with equivalent amount of DMSO. Cells were then further processed for Mass Spectrometry (MS), WB or immunofluorescence (IF) analysis.

### Isolation of IACs

IACs were isolated from cells cultivated in 10 cm diameter Petri dishes for 48 hours post-transfection for MDA-MB-435S cells or 72 hours post-transfection for RPMI-7951, with or without CytoD treatment, as described previously (Loncaric et al., 2023, Paradzik et al., 2020, Jones et al., 2015). Briefly, cells were washed with DMEM-HEPES and incubated with Wang and Richard’s reagent (DTBP, 6 mM, Thermo Fisher Scientific) for 10 minutes (MDA-MB-435S) or 15 minutes (RPMI-7951). DTBP was quenched with 0.03M Tris-HCl (pH 8) and cells were lysed using modified RIPA buffer (50 mM Tris-HCl, pH 7.6; 150 mM NaCl; 5 mM disodium EDTA, pH 8; 1% (w/v) Triton X-100, 0.5% (w/v) SDS, 1% (w/v) sodium deoxycholate). Cell bodies were removed by high-pressure washing with tap water for 10 seconds, and remaining IACs were collected by scraping into adhesion recovery solution (125 mM Tris-HCl, pH 6.8; 1% (w/v) SDS; 150 mM dithiothreitol). Samples containing isolated IACs were acetone-precipitated and further processed for MS or WB analysis (Robertson et al., 2015).

### Sample Preparation for MS, Data Analysis, Protein-Protein Interaction Network Analysis and Functional Enrichment Analysis

For MS analysis, isolated IAC samples were prepared using in-gel trypsin digestion (Jones et al., 2015, Paradzik et al., 2020), and analysed using a modified version of the LC-MS/MS method, as previously described (Horton et al., 2015). Samples were analysed by LC-MS/MS using an UltiMate 3000 Rapid Separation LC (RSLC, United States) coupled to an Orbitrap Exploris mass detector (Thermo Fisher Scientific, United States) with electrospray ionization. Peptide mixtures were eluted for 60 minutes. To identify proteins after MS analysis, data were searched against the human Swissprot and Trembl database (July 2022, July 2023) using Mascot (Matrix science, version 2.5.1). Fragment ion tolerance was set to 0.6 Da, while parent ion tolerance was 10 PPM. We set the carbamidomethylation of cysteine as a fixed modification and oxidation of methionine as a variable modification. In further analysis, we considered only peptides with ions of charge precursors 2+, 3+ and 4+. Protein identifications were further refined using Scaffold (Proteome Software, version 5.1.0). Protein (99%) and peptide (90%) probabilities were assigned using the Protein Prophet algorithm (Nesvizhskii et al., 2003) as incorporated by Scaffold including a minimum of four spectral counts per protein. Spectral counts were used as a measure of protein abundance. QSpec statistical method (Choi et al., 2008) was used for MS data to measure the significance of differentially identified proteins in control cells and treated cells. For visualization of differentially expressed proteins, volcano plot (GraphPad Prism version 9.0.0 (GraphPad Software)) with the following setup was created: −Log (FDR) ≥ 0.05 corresponding to FDR ≤ 0.05; and fold change ≥ 1.5 or 2.

### SDS-PAGE and Western Blotting

Total cell lysates were obtained from 3.5 cm Petri dishes in 200 µL RIPA buffer supplemented with protease inhibitor cocktail (ThermoFisher). Samples for SDS-PAGE were collected by scraping. Samples containing an equal amount of protein were mixed in 6× Laemmli loading buffer (375 mM Tris-HCl (pH 6.8), 30% (w/v) glycerol, 12% (w/v) SDS, 0.02% (w/v) bromophenol blue, 12% (v/v) 2-mercaptoethanol) to reach a final 1× concentration, sonicated and heated for 5 minutes at 96°C. Isolated IACs were prepared for SDS-PAGE by solubilization in 2× Laemmli loading buffer and heating for 20 minutes at 70°C while shaking (1000 rpm). All samples were loaded onto pre-casted gradient gel (4 – 15% Mini-PROTEAN TGX) (Bio-Rad), separated by SDS-PAGE and semi-dry transferred to nitrocellulose (Bio-Rad). The membrane was blocked in 5% (w/v) non-fat dry milk or 5% (w/v) bovine serum albumin (BSA, Carl Roth), and incubated overnight with the appropriate antibodies, followed by incubation with horseradish peroxidase coupled secondary antibody. The primary and secondary antibodies are listed in Additional file 1 Table S1. Detection was performed using chemiluminescence (PerkinElmer) and documented with Uvitec Alliance Q9 mini (BioSPX b.v.). Blots were quantified using ImageJ.

### Confocal Microscopy

For IF analysis, cells were plated on coverslips in 24-well plates. After 48 hours, cells were fixed using ice-cold methanol for 10 minutes or 2% (v/v) paraformaldehyde for 12 minutes followed by permeabilization with 0.1% (v/v) Triton X-100 for 2 minutes, and stained with protein specific primary antibodies for 1 hour, followed by incubation with conjugated secondary antibodies for 1 hour. The primary and secondary antibodies are listed in Additional file 1 Table S1. Actin stress fibers were stained with rhodamine phalloidin (Cell Signaling Technology). Cells were mounted with DAPI Fluoromount-G (SouthernBiotech) and fluorescence and respective IRM images were acquired using an inverted confocal microscope (Leica TCS SP8 X, Leica Microsystems) with the HC PL APOCS2 63×/1.40 oil-immersion objective, zoom set at 2×. Images were analysed using LAS X software 3.1.1 (Leica Microsystems) and ImageJ. All images were taken with the focus adjusted to the adhesion sites of cells at the upper surface of glass coverslip.

### Proximity ligation assay

Proximity ligation assay (PLA), using Duolink® PLA technology (Alam, 2018), was performed according to manufacturer’s instructions. Briefly, cells seeded on coverslips for 48 hours were fixed, blocked and incubated with selected primary antibodies for 1 hour at 37°C in a preheated humidity chamber. Primary antibodies are listed in Additional file 1 Table S1. Following washing coverslips were incubated with secondary antibodies conjugated to proprietary oligonucleotide arms (Navenibodies) for 1 hour. Incubation with specific enzymes enables the formation and amplification of a DNA circle in the spots of protein proximity. Fluorescent dots, generated by binding of fluorescently labelled probes, were visualized using an inverted confocal microscope (Leica TCS SP8 X, Leica Microsystems) with the HC PL APOCS2 63×/1.40 oil-immersion objective, zoom set at 2×. Images were analysed using LAS X software 3.1.1 (Leica Microsystems) and ImageJ.

## RESULTS

### Both melanoma cell lines MDA-MB-435S and RPMI-7951 display integrin αVβ5-positive, vinculin-negative structures in the cell centre

RAs primarily require integrin αVβ5, lack association with actin, and are devoid of markers of FAs, including vinculin (Lock et al., 2018, Zuidema et al., 2018). We have previously analysed IACs of two melanoma cell lines MDA-MB-435S and RPMI-7951, grown in long-term culture, using biochemical isolation and MS-based proteomics, and demonstrated that both cell lines preferentially use integrin αVβ5 for adhesion (Paradzik et al., 2020, Stojanović et al., 2025). These adhesome datasets were obtained from cells cultured for 48 hours (MDA-MB-435S) or 72 hours (RPMI-7951), on culture plates without prior ECM coating, to allow the cells to regulate their own environment by secreting ECM proteins. In these cultures, integrin αVβ5 was localized at the periphery in vinculin-positive FAs, but also in the central part of the cell in vinculin-negative structures, which were not associated with actin fibers (Figure 1). This suggests that both melanoma cell lines, MDA-MB-435S and RPMI-7951, form RAs.

**Figure 1.**
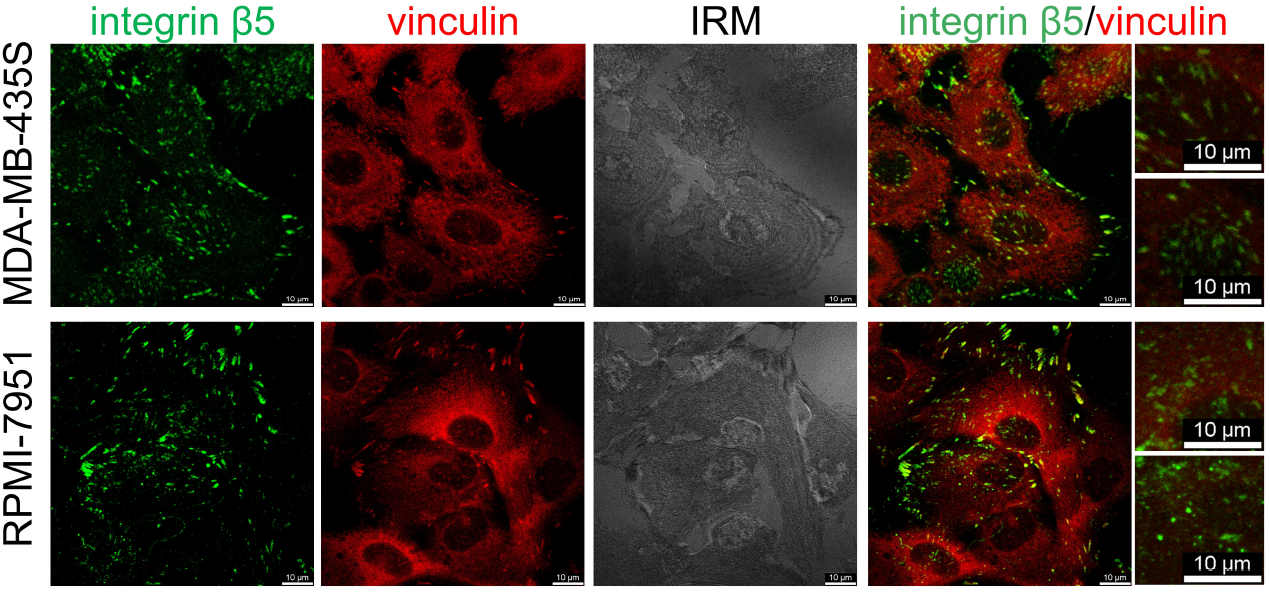
Melanoma cell lines MDA-MB-435S and RPMI-7951 contain integrin β5-positive, vinculin negative structures in the cell centre. Forty-eight hours after seeding, cells were fixed with 4% PFA, and stained with anti-vinculin antibody followed by Alexa-Fluor IgG1 555-conjugated antibody (red) and anti-integrin β5 antibody followed by Alexa-Fluor IgG2b 488-conjugated antibody (green). Interference reflection microscopy (IRM) images were taken. Analysis was performed using TCS SP8 Leica. Scale bar = 10 μm.

### Composition of IACs not associated with actin in melanoma cell lines MDA-MB-435S and RPMI-7951

Since RAs are formed independently of the actin cytoskeleton both cell lines were exposed to CytoD, an inhibitor of actin polymerisation known to disrupt FAs, as previously described (Lock et al., 2018, Zuidema et al., 2022). To determine the optimal CytoD concentration for enriching RAs, cells were treated with increasing doses for 2 hours and IF was employed to visualize F-actin, the FA marker vinculin, the RA component Numb and the nucleus (Lock et al., 2018). CytoD disrupted actin and FAs in a dose-dependent manner. Numb remained unaffected and progressively clustered beneath the nucleus, in a pattern characteristic of RAs. The most pronounced effect was observed at the CytoD concentration of 20 μM, as described in (Lock et al., 2018) and was used for all subsequent experiments (Suppl. Figure S1).

Next, IACs were isolated using a previously optimized protocol (Paradzik et al., 2020, Stojanović et al., 2025) from CytoD treated cells with enriched RAs (CytoD(+), i.e. RAs) and compared to IAC isolates from cells treated with solvent DMSO (CytoD(-), i.e. FAs and RAs). IAC isolation and LC-MS/MS analysis were performed from four biological replicates of MDA-MB-435S (Table S2.1) and RPMI-7951 (Table S2.2) cells, and spectral counts were used as a measure of protein abundance. MS analysis identified 746 proteins for MDA-MB-435S and 655 proteins for RPMI-7951 cells.

Integrin subunits that appeared in CytoD(-) samples of both cell lines were αV, β5 and β1 (Table S2.1, S2.2). The most abundant integrin subunits were αV and β5, again confirming previously published data that both cell lines preferentially use integrin αVβ5 for adhesion (Paradzik et al., 2020, Loncaric et al., 2023, Stojanović et al., 2025).

To determine the differences in IAC composition between CytoD(-) and CytoD(+) samples, in both cell lines, the method of Lock et al. (Lock et al., 2018) was employed. They defined proteins as being RA components if a greater than 2-fold increase in abundance was observed following CytoD treatment. This threshold was lowered to 1.1 since in our experiments these ratios for typical RA proteins were lower. Another reason for using a lower ratio was to enable the detection of proteins present in both FAs and RAs, such as talin2. This narrowed the list of candidate proteins to 225 for MDA-MB-435S cells and 430 for RPMI-7951 cells (Table S2.1., S2.2.), of which 98 were shared between the cell lines (Figure 2A, Table S2.3.) and became the focus of future analysis. Stonin1, which has been identified as a marker for RAs (Lukas et al., 2024) was identified with a low number of spectra only in MS samples of MDA-MB-435S, but not RPMI-7951 cells. This could be due to the fact that stonin1 exhibits variable expression across different human cell types (Lukas et al., 2024).

**Figure 2.**
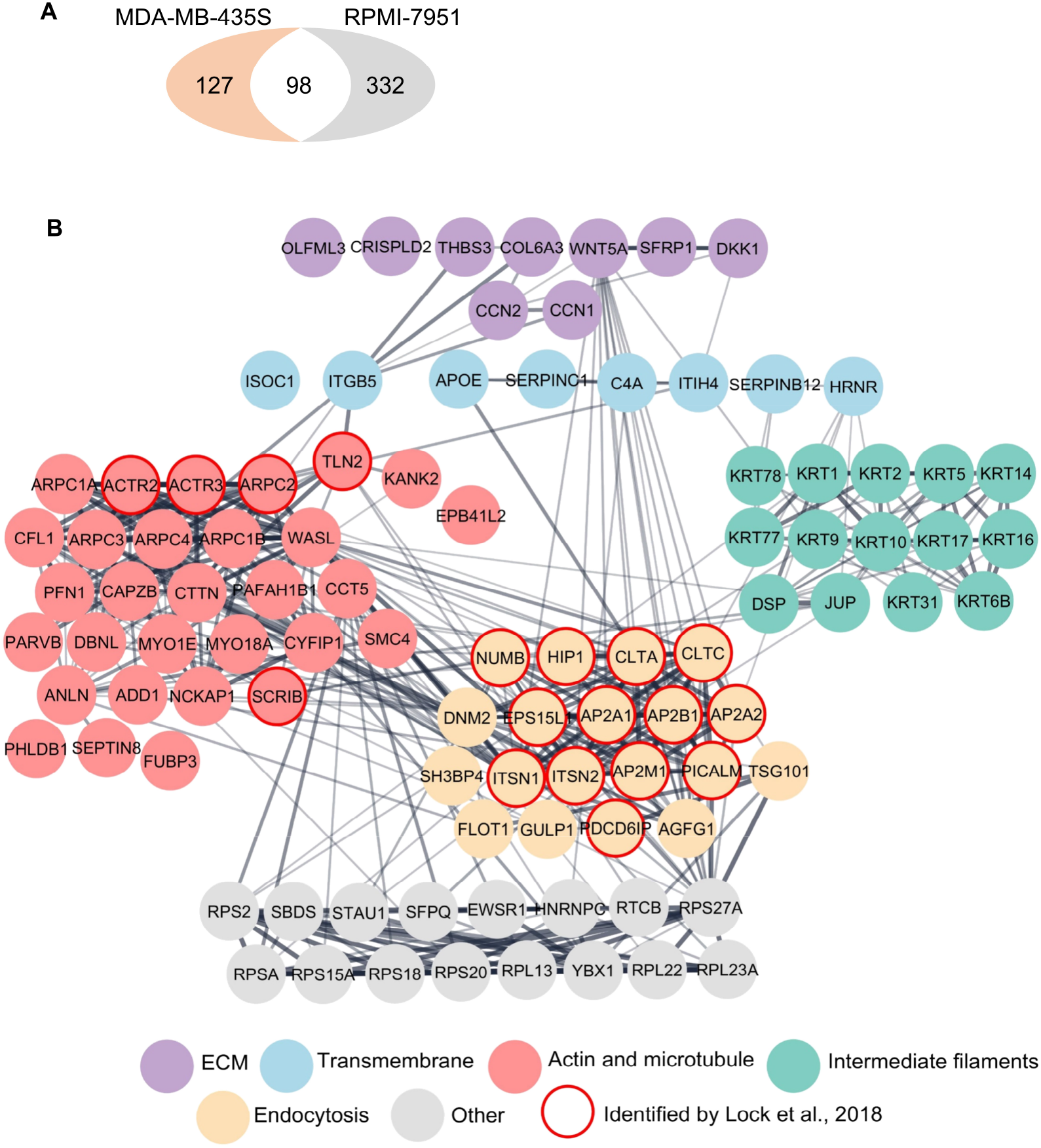
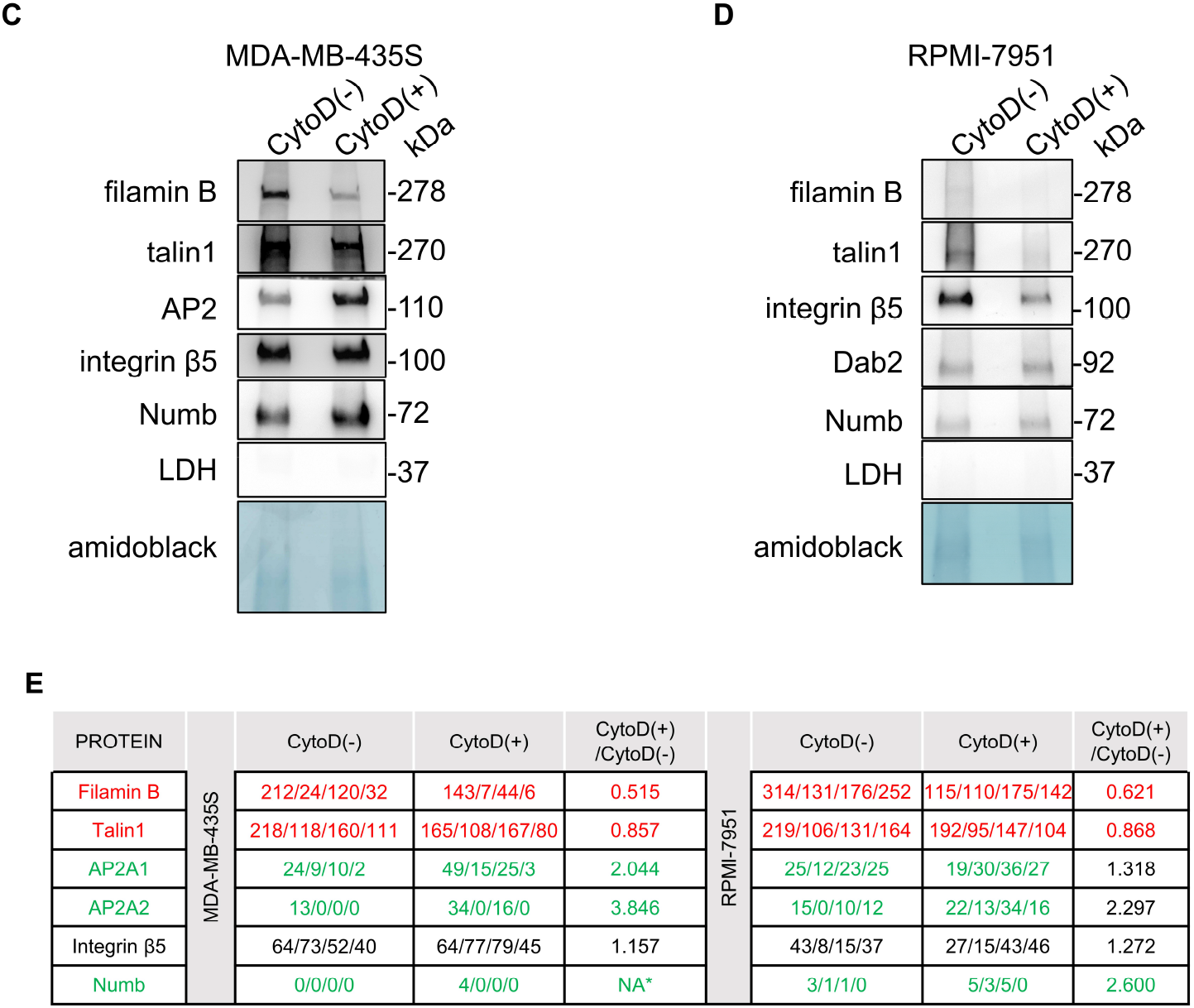
MS analysis of RAs isolated from MDA-MB-435S and RPMI-7951 melanoma cell lines. (A) Venn diagram – number of RA proteins enriched only in MDA-MB-435S cells (orange) and number of RA proteins enriched only in RPMI-7951 cells (grey). Enriched RA proteins found in both cell lines shown in the intersected white area of the diagram. (B) PPI network identified by MS in RAs isolated from both MDA-MB-435S and RPMI-7951 cells. Identified proteins are labelled with gene symbols, arranged and coloured according to their functional group as indicated. (C, D) MS data validation. WB analysis of IAC proteins in MDA-MB-435S cells (C) and RPMI-7951 cells (D) and comparison of non-treated (CytoD(-)) and CytoD treated (CytoD(+)) samples. Two hours upon CytoD treatment, IACs were isolated and WB analysis was performed. Amidoblack staining of the membrane was used as a loading control. The results presented are representative of two independent experiments yielding similar results. (E) Exempt from MS data (Table S2.1 and S2.2.) showing number of protein-specific spectra validated in (C, D). Dataset consists of four replicas. CytoD(+)/CytoD(-) ratio shown.

A constructed Protein-Protein Interaction Network (PPI) network of the 98 enriched proteins (Figure 2B; Table S2.1, S2.2.) included several ECM proteins including collagen type VI alpha 3 chain (COL6A3), cysteine-rich angiogenic inducer 61 CYR61, i.e. CCN1), thrombospondin-3 (THBS3) and Wnt family member 5A (WNT5A). Transmembrane proteins identified within the network included integrin αV (ITGAV) and β5 (ITGB5), apolipoptotein E (APOE), and hornerin (HRNR).

Actin and microtubule-related proteins identified within the PPI network (Figure 2B) comprised actin-related proteins 2 and 3 (ACTR2, ACTR3), components of the Arp2/3 complex (ARPC1A, ARPC1B, ARPC2, ARPC3, ARPC4), myosin IE and myosin XVIIIA (MYO1E, MYO18A), cortactin (CTTN), anillin (ANLN), talin2 (TLN2), cofilin-1 (CFL1), septin-8 (SEPTIN8) and others. It has been shown that Arp2/3-mediated actin polymerization through cortactin (Uruno et al., 2001) facilitates the transition of clathrin plaques into endocytic pits, underscoring a mechanistic role for actin in endocytosis at plaque peripheries (Leyton-Puig et al., 2017). In addition to talin2, known to be present in FAs and RAs (Lock et al., 2018), MS data also showed slightly increased abundance of KANK2 in MDA-MB-435S cells (Table S2.1, S2.2., S2.3; Figure 3A), which is a known interactor of talin2 (Loncaric et al., 2023). Although increased abundance of KANK2 was observed in RPMI-7951 cells, the number of spectra was lower (Table S2.1, S2.2., S2.3; Figure 3A). Nevertheless, this increased abundance of KANK2 indicated that KANK2 might be a constituent of RAs.

**Figure 3.**
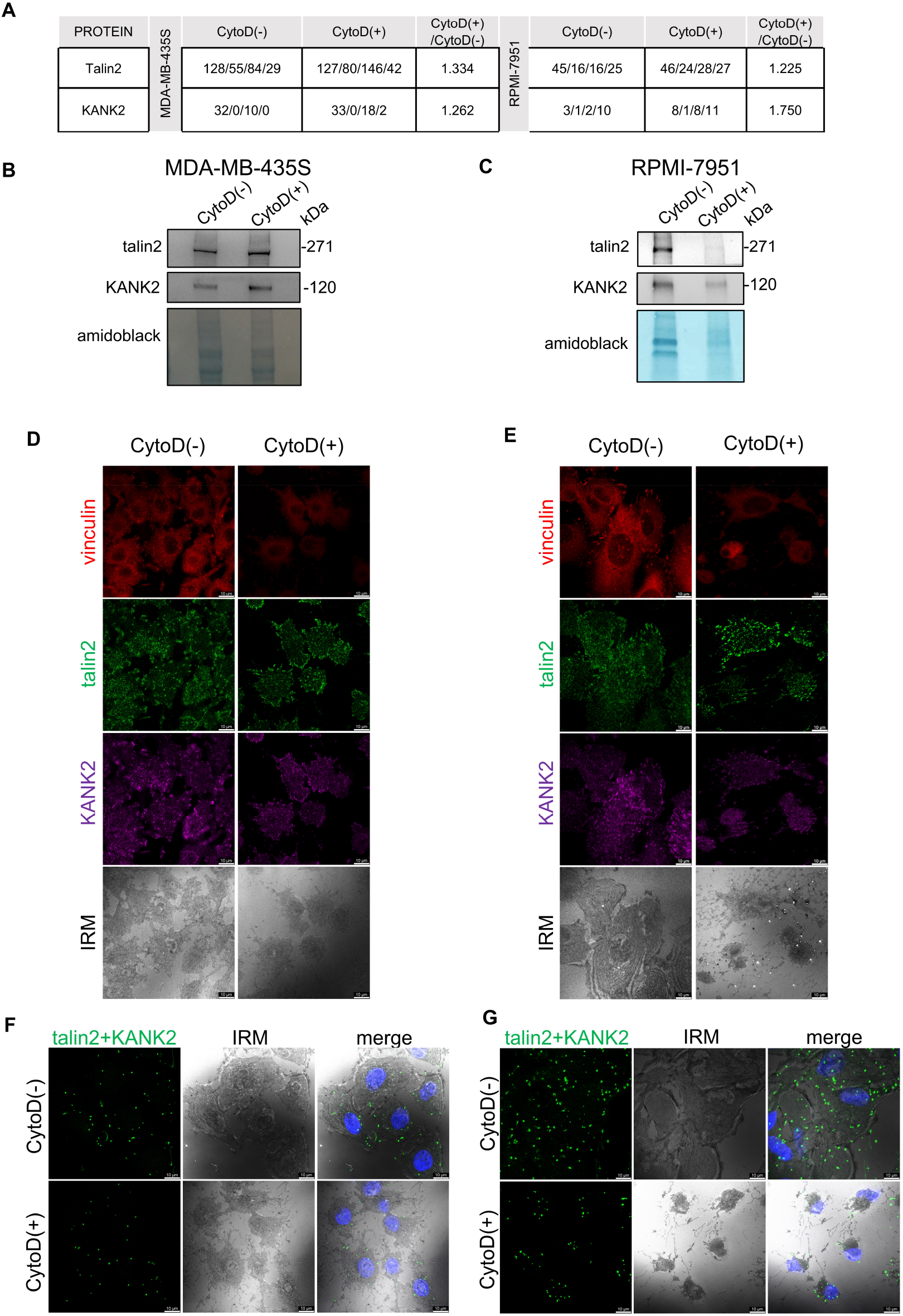
KANK2 colocalizes with talin2 in vinculine-negative structures in two melanoma cell lines MDA-MB-435S and RPMI-7951. (A) MS data of number of spectra specific for talin2 and KANK2 in MDA-MB-435S and RPMI-7951 cells. Dataset consists of four replicas. CytoD(+)/CytoD(-) ratio shown for both cell lines. (B, C) MS data validation. WB analysis of talin2 and KANK2 in MDA-MB-435S (B) and RPMI-7951 cells (C) in non-treated (CytoD(-)) and CytoD treated (CytoD(+)) samples. Two hours upon CytoD treatment, IACs were isolated and WB analysis was performed. Amidoblack staining of the membrane was used as a loading control. The results presented are representative of two independent experiments yielding similar results. (D, E) Talin2 and KANK2 are localized within RAs of both cell lines. Forty-eight hours after seeding, and 2 hours after CytoD treatment, cells were methanol fixed, stained with anti-vinculin antibody followed by Alexa-Fluor IgG1 555-conjugated antibody (red), anti-talin2 antibody followed by Alexa-Fluor IgG2b 488-conjugated antibody (green) and anti-KANK2 antibody followed by Alexa-Fluor 647-conjugated antibody (magenta). IRM images were taken. Analysis was performed using TCS SP8 Leica. Scale bar = 10 μm (F, G) Talin2 and KANK2 are in close proximity in RAs of both cell lines. Forty-eight hours after seeding, and 2 hours after CytoD treatment, RPMI-7951 cells were methanol fixed, and a PLA assay with anti-talin2 and anti-KANK2 primary antibody was performed. Generated fluorescent dots were visualized and interference reflection microscopy (IRM) images were taken. Analysis was performed using TCS SP8 Leica. Scale bar = 10 μm.

There was also an increase in abundance of several keratins in both cell lines (Figure 2B). Since keratins can originate from MS sample preparation contamination (Hodge et al., 2013) and provide false positives, and thus far, no literature data link keratins to RAs, we did not evaluate their significance.

CytoD(+) samples in MDA-MB-435S and RPMI-7951 cells showed a higher abundance of clathrin adaptor proteins AP2A1, AP2A2, AP2B1 and AP2M1, clathrin light and heavy chains CLTA and CLTC, Numb, phosphatidylinositol-binding clathrin assembly protein (PICALM), PTB domain-containing engulfment adaptor protein 1 (GULP1, programmed cell death 6-interacting protein (PDCD6IP), SH3 domain-binding protein 4 (SH3BP4), Huntingtin-interacting protein 1 (HIP1), flotillin 1 (FLOT1), Arf-GAP domain and FG repeat-containing protein 1 (AGFG1), scaffold proteins in endocytosis intersectins 1 and 2 (ITSN1 and ITSN2), dynamin2 (DNM2) and epidermal growth factor receptor substrate 15-like 1 protein (EPS15L1) (Figure 2B). It should be noted that, of the 98 proteins enriched upon CytoD treatment, 18 are part of the reticular adhesome determined by Lock et al. (Lock et al., 2018) (Figure 2B (circled red), Table S2.4.), and 13 of those are proteins involved in endocytosis, while the rest are actin-related.

MS analysis identified several ribosomal proteins (ribosomal protein large (RPL) and small (RPS)) and those related to RNA and protein synthesis such as ribosome maturation factor SBDS, RNA 2’,3’-cyclic phosphate and 5’-OH ligase RTCB and proline- and glutamine-rich splicing factor SFPQ. Previous studies have described the presence of ribosomes in FAs and their transport towards adhesion sites upon binding to the ECM (Simpson et al., 2020). Ribosomal protein SA (RPSA), first identified as a laminin-binding protein, has been shown to regulate ribosome biogenesis, cytoskeletal organisation, and nuclear functions (Mercurio, 1995, DiGiacomo and Meruelo, 2016). It has also been implicated in cell migration (Lefebvre et al., 2020), tumor cell blebbing and extracellular vesicle shedding (Brassart et al., 2019). However, the role of these proteins in RA-enriched samples remains unknown.

We validated MS results using WB analysis (Figure 2C, D, E, Suppl. Figure S3, S4). A decrease in abundance of filamin B and talin1 (in both cell lines) in the CytoD(+) compared to the CytoD(-) samples confirmed the loss of FAs. Successful isolation of RAs was further demonstrated by increased AP2 and Numb (in MDA-MB-435S), and Numb and Dab2 levels (in RPMI-7951) in the CytoD(+) samples, which is in line with the literature (Lock et al., 2018, Zuidema et al., 2022, Zuidema et al., 2018). Integrin β5 was expressed in all samples (Figure 2C, D, E, Suppl. Figure S3, S4), consistent with its previously confirmed localization in both FAs and RAs (Lock et al., 2018, Zuidema et al., 2018). In conclusion, the results demonstrate that both MDA-MB-435S and RPMI-7951 cells form αVβ5 RAs.

### KANK2 is a component of RAs in MDA-MB-435S and RPMI-7951 cells

In addition to proteins we expected to be maintained or enriched in CytoD(+) samples based on the literature (Lock et al., 2018), we observed an enrichment in KANK2 following CytoD treatment in both MDA-MB-435S and RPMI-7951 cells (Figure 3A, B, C, Suppl. Figure S5, S6), a protein not previously recognized as a component of RA. WB analysis shows KANK2 presence in CytoD(+) samples from both MDA-MB-435S and RPMI-7951 cells confirming that KANK2 is present in RAs (Figure 3B, C, Suppl. Figure S5, S6). IF analysis demonstrated that talin2, which we recently showed to colocalize with KANK2 in both FAs and FBs (Loncaric et al., 2023, Stojanović et al., 2025), also colocalized with KANK2 upon CytoD treatment, when FAs were disrupted (indicated by the loss of FA marker vinculin) (Figure 3 D,E). This indicates that the colocalizing talin2 and KANK2 are components of RAs. Finally, PLA following CytoD-induced actin disruption further confirmed the proximity of KANK2 and talin2 in RAs (Figure 3F, G).

Alongside the integrin receptor subunits αV, β5 and β1, the adhesome of RPMI-7951 cells also contained integrin α5, though at low levels (Table S2.2). Our recent data show that RPMI-7951 cells form FBs, containing talin2 and KANK2 (Stojanović et al., 2025). In contrast, the adhesome of MDA-MB-435S cells contained integrin β1 alongside αV and β5, but there were no spectra for integrin α5 (Table 2.1). Under conditions where cells were seeded on an uncoated surface and cultured, the elongated, integrin α5-positive structures characteristic of FBs were not observed (Supplementary Fig. S2), but these were evident in RPMI-7951 cells (Stojanović et al., 2025). To additionally confirm that KANK2 and talin2 are indeed components of RAs, cells were transfected with integrin subunit α5-specific siRNA (to reduce FBs), treated them with CytoD (to disrupt FAs), and PLA performed. This revealed a talin2-KANK2 positive signal in both cell lines (Fig 4). Based on these results, KANK2 is demonstrated to be a hitherto unidentified component of RAs.

**Figure 4.**
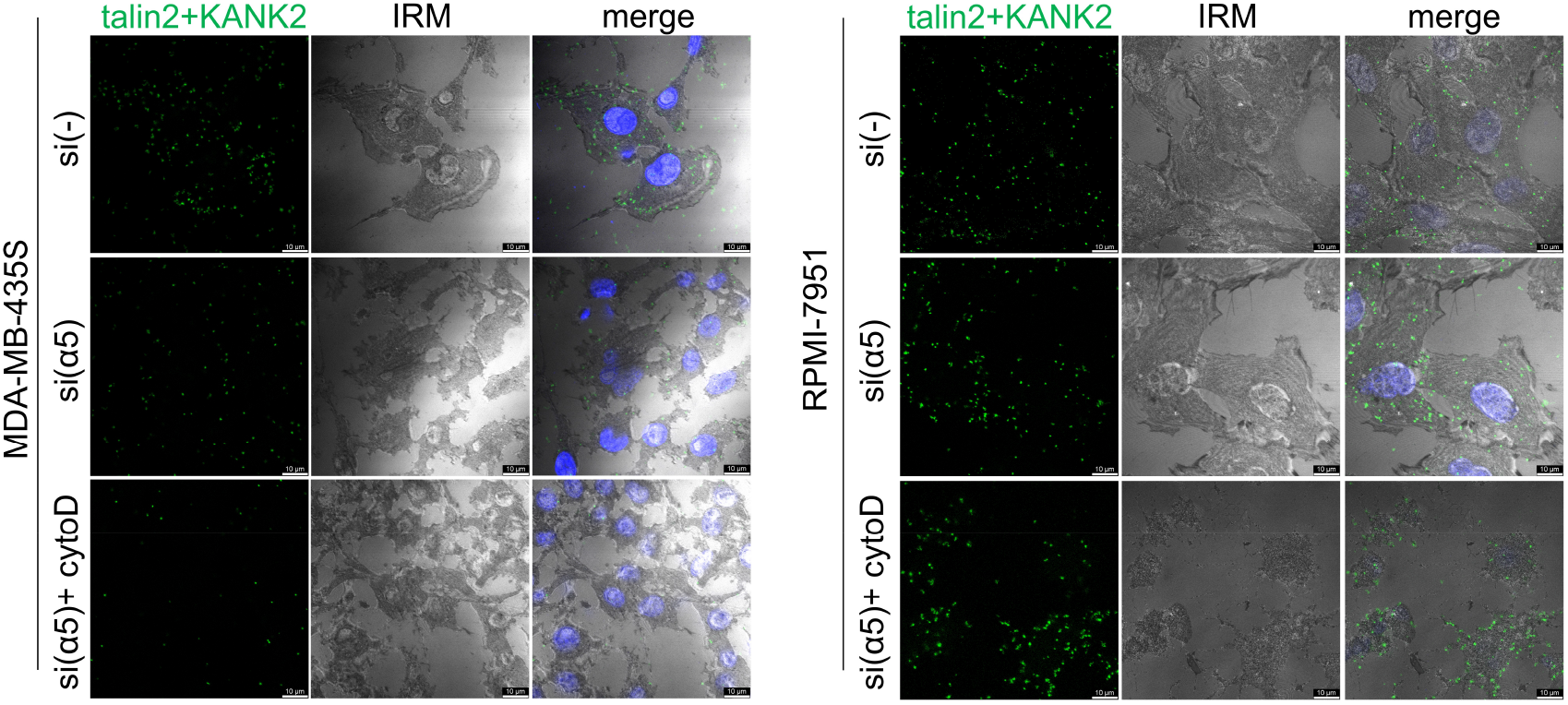
KANK2 is a component of RAs in two melanoma cell lines. Talin2 and KANK2 are in close proximity in RAs of both cell lines. Forty-eight hours after transfection with either control siRNA (si(-)) or integrin α5-specific siRNA (si(ITGA5)), followed by 2 hours CytoD treatment, cells were methanol fixed, and a PLA assay with anti-talin2 and anti-KANK2 primary antibody was performed. Generated fluorescent dots were visualized and IRM images were taken. Analysis was performed using TCS SP8 Leica. Scale bar = 10 μm.

### The effect of talin2 or KANK2 knockdown on RAs abundance in MDA-MB-435S and RPMI-7951 cells

In MDA-MB-435S cells, talin2 and KANK2 within integrin αVβ5 FAs functionally interact to regulate microtubule dynamics and cell migration. More specifically, knockdown of either talin2 or KANK2 mimics the effect of integrin αV or β5 subunit knockdown, resulting in reduced cell migration (Loncaric et al., 2023). However, in RPMI-7951 cells, talin2 and KANK2 colocalize in FAs and FBs (Stojanović et al., 2025), and their knockdown has differential effects on cell migration. Talin2 knockdown mimicked the effect of αV or β5 integrin knockdown i.e. reduced cell migration, while KANK2 knockdown resembled the effect of integrin α5 knockdown, leading to increased cell migration. Specifically, KANK2 knockdown increased the size of α5-positive, but not β5-positive, structures. These findings suggest that the localization of KANK2 to FAs or FBs differentially influences integrin αVβ5 FAs and migration in melanoma cell lines (Stojanović et al., 2025).

To investigate how talin2 or KANK2 knockdown affects their mutual localization and/or abundance in RAs, each protein was knocked down in MDA-MB-435S and RPMI-7951 cells, exposed them to CytoD, and their localization analysed using IF. The loss of vinculin signal verified CytoD treatment effectiveness. In both cell lines, talin2 and KANK2 knockdowns had no effect on either signal in RAs (Figure 5). Therefore, neither talin2 nor KANK2 are essential for RA maintenance.

**Figure 5.**
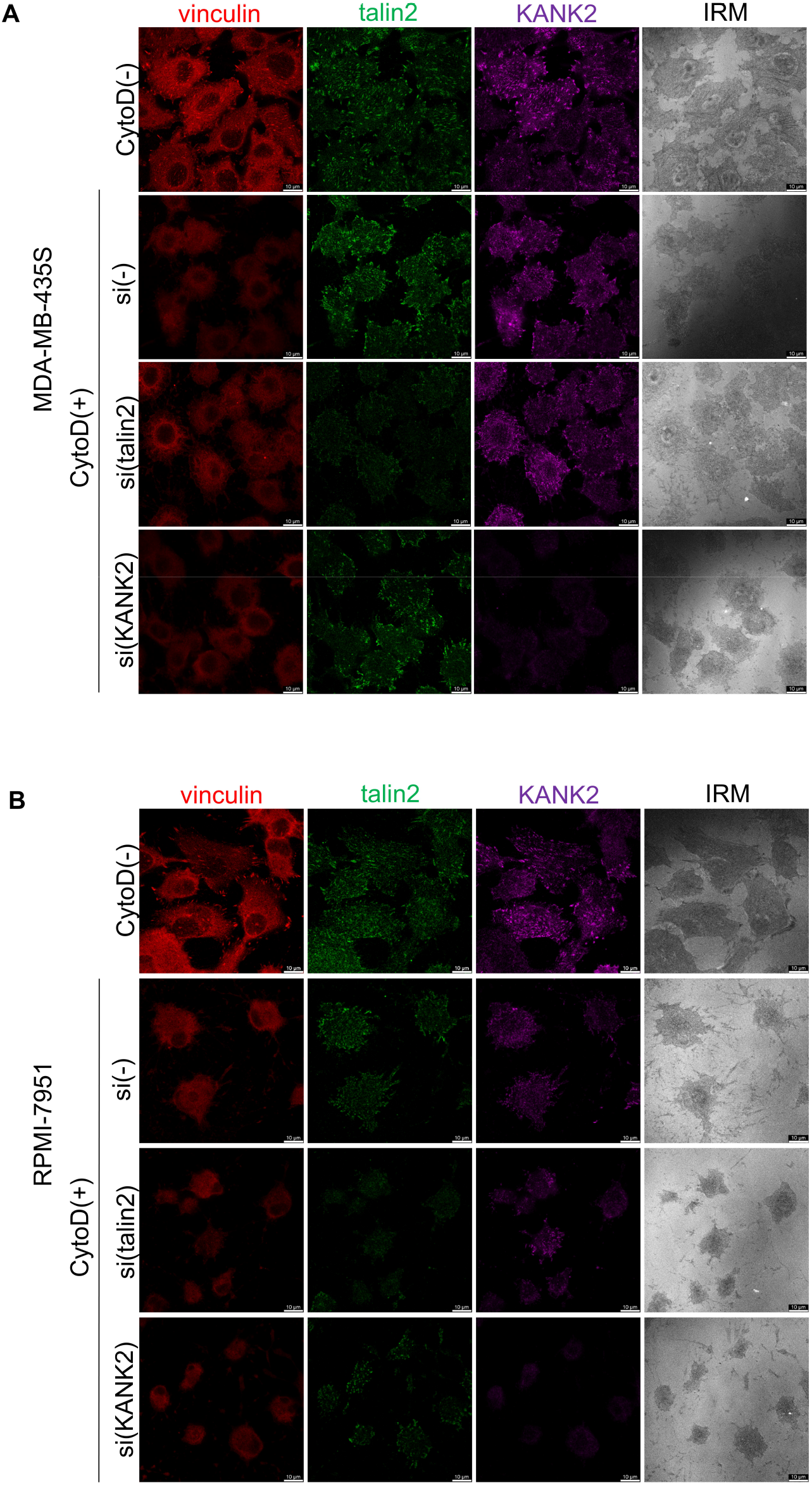
Neither talin2 nor KANK2 are essential for RA maintenance. Forty-eight hours after transfection with either control siRNA (si(-)), talin2-specific (si(talin2)) or KANK2-specific (si(KANK2)) siRNA, followed by 2 hours CytoD treatment, MDA-MB-435S (A) and RPMI-7951 (B) cells were methanol fixed, and stained with anti-vinculin antibody followed by Alexa-Fluor IgG1 555-conjugated antibody (red), anti-talin2 antibody followed by Alexa-Fluor IgG2b 488-conjugated antibody (green) and anti-KANK2 antibody followed by Alexa-Fluor 647-conjugated antibody (magenta). IRM images were taken. Analysis was performed using TCS SP8 Leica. Scale bar = 10 μm.

The next step was to analyse RA composition and abundance following talin2 or KANK2 knockdown. For MDA-MB-435S cells, three biological replicates, while for RPMI-7951 two biological replicates per sample were analysed. MS results (Table S3.1, S3.1.1, S3.1.2, S3.2, S3.2.1, S3.2.2) were processed, analysed as described above, and visualized using volcano plots. In both cell lines, successful knockdown of talin2 and KANK2 was confirmed by a reduced number of spectra for talin2 and KANK2, respectively (Figure 6).

**Figure 6.**
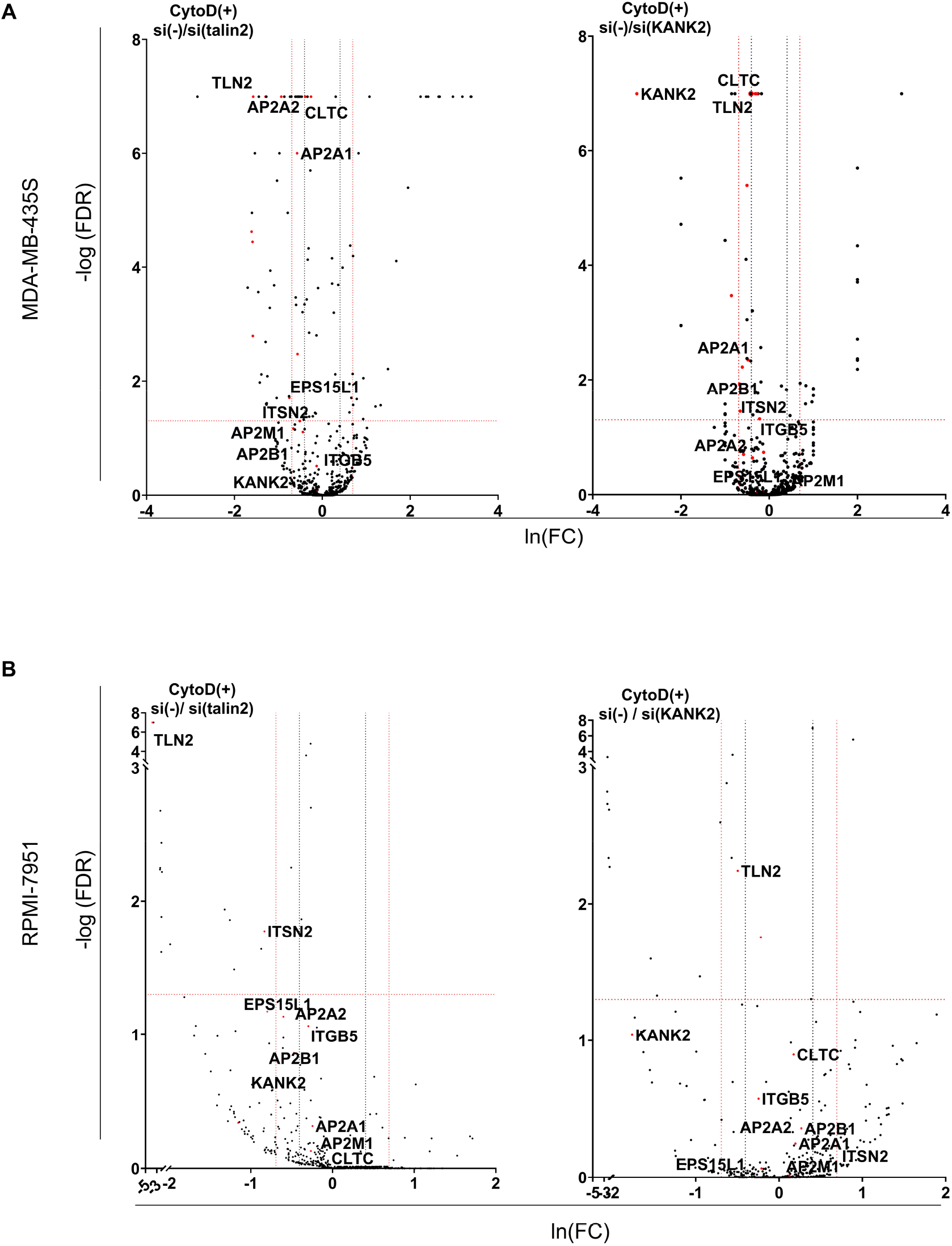
Talin 2 and KANK2 knockdown affect the abundance of RA components differently in MDA-MB-435S than in RPMI-7951 cells. Volcano plot analysis of proteins detected in RAs isolated from (A) MDA-MB-435S and (B) RPMI-7951 cells transiently transfected with control siRNA (si(-)) versus talin2-specific siRNA (si(talin2)) or KANK2-specific siRNA (si(KANK2)). RA proteins are visualized as volcano plot after the analysis with QSpec/QProt to generate -Log (FDR) and fold change values. Cut off values of −Log (FDR) ≥ 0.05 (red horizontal dotted line) corresponding to FDR ≤ 0.05; and fold change ≥ 1.5 (black vertical dotted line) or 2 (red vertical dotted line) were used. Each dot on the plot represents one protein. Proteins with significantly different abundance between RAs of si(talin2) or si(KANK2) and si(-) cells and of interest for this paper were marked with their gene name. Upper left quadrant – proteins detected with lower levels of spectra in si(talin2), upper right quadrant – proteins detected with higher levels of spectra in si(talin2) or si(KANK2).

In MDA-MD-435S cells, talin2 knockdown did not affect KANK2 levels, while in RPMI-7951 cells, the very low number of KANK2-specific spectra made it impossible to determine whether KANK2 levels changed following talin2 knockdown. Conversely, in both cell lines, KANK2 knockdown reduced talin2 levels. These data indicate the dependence of talin2 on KANK2 (Figure 6; Table S3.1, S3.1.1, S3.1.2, S3.2, S3.2.1, S3.2.2). However, since RA formation is independent of talin2 (Lock et al., 2018), it is very likely to be independent of KANK2.

Next, the effect of knockdown of talin2 or KANK2 on the abundance of RA components was examined. A significant change in the abundance of characteristic RA proteins was observed following talin2 knockdown in both cell lines, and after KANK2 knockdown in MDA-MB-435S, but not in the RPMI-7951 cell line (Figure 6). In MDA-MB-435S cells, there was a significant reduction in the amount of several RA components such as AP2A1, AP2A2, EPS1L15, and ITSN2. KANK2 knockdown in MDA-MB-435S cells also led to a decrease in the level of several RA components such as CTLC proteins, AP2A1, AP2B1, and ITSN2 (Figure 6A). In RPMI-7951 cells, knockdown of talin2 similarly led to a decrease in the level of RA components AP2A1, AP2A2, AP2B1, CTLC, EPS1L15 and ITSN2 (Figure 6B). These results are in line with observed changes in FAs of MDA-MB-435S after either talin2 or KANK2 knockdown and RPMI-7951 cells after talin2 knockdown. Thus, FAs become less dynamic and their size increases but not their number, indicating the retention of FA proteins (Loncaric et al., 2023, Stojanović et al., 2025), which leads to a reduction in RAs.

KANK2 knockdown did not have the same effect as talin2 knockdown in RPMI-7951 cells. The level of the RA components AP2A1, AP2A2, AP2B1, CTLC, EPS1L15 and ITSN2 did not change (Figure 6B). This is in line to what happens with integrin αVβ5 FAs after KANK2 knockdown. Namely, KANK2 knockdown in RPMI-7951 did not affect integrin β5-positive adhesions at all (Stojanović et al., 2025).

## DISCUSSION

Here, we report the composition of RAs in two melanoma cell lines, MDA-MB-435S and RPMI-7951, and compare the identified components with previously published data. The findings confirm previously obtained data by Lock et al. (Lock et al., 2018) that RAs contain talin2. Additionally, KANK2 is shown to be a novel component of RAs. Given that we have extensively studied the role and localization of talin2 and KANK2 in FAs and FBs of these two cell lines (Loncaric et al., 2023, Stojanović et al., 2025), we further analysed how the knockdown of each protein affects RAs. Our results demonstrate that talin2 and KANK2 are not essential for the formation of RAs; however, their knockdown influences RA component abundance, and primarily mirrors the roles of talin2 and KANK2 in αVβ5 FAs. This highlights the importance of adhesion crosstalk in cellular function.

The analysis of IACs in the melanoma cell lines MDA-MB-435S and RPMI-7951 demonstrated that both cell lines in long-term culture preferentially use integrin αVβ5 for adhesion, forming FAs (Loncaric et al., 2023, Stojanović et al., 2025) and RAs (this work) that contain talin2 and KANK2. It was also found that, unlike MDA-MB-435S cells, RPMI-7951 cells form FBs that also contain talin2 and KANK2 (Stojanović et al., 2025). Here RAs were analysed in both cell lines and it was demonstrated that clathrin and endocytic adaptor proteins are enriched in RA isolates as previously reported by others (Lock et al., 2018, Zuidema et al., 2022, Zuidema et al., 2018). In the analysis of RA proteins, the same logic as Lock and colleagues (Lock et al., 2018) was followed; i.e. the proteins we considered to be potential RA proteins were those maintained or enriched in the sample during RA isolation. In doing so, the presence of most of the previously identified proteins was confirmed in the melanoma cells (Lock et al., 2018). Stonin1, which has been identified as a specific adaptor for the endocytosis of NG2 and as an important factor for FA dynamics and cell migration (Feutlinske et al., 2015), and recently shown to be as a marker for RAs (Lukas et al., 2024), was only detected in RA isolates of MDA-MB-435S, with a low number of spectra. In RPMI-7951 RA isolates, stonin1 was not detected, nor was it detected in RPMI-7951 IAC isolates (Stojanović et al., 2025). This could be a consequence of the protein not being expressed in abundance in the melanoma cells. Therefore, in subsequent analyses of RA levels, endocytic adaptors such as AP2 as markers of RAs were used as markers.

Previous research identified integrin αVβ5 as the main receptor in RAs; however, it is still not clear whether β3 and β1 can also contribute to their formation (Zuidema et al., 2018, Lock et al., 2018). MS analysis of isolated IACs after disruption of actin cytoskeleton using CytoD confirmed that αVβ5 integrin preferentially forms RAs in the two melanoma cell lines. Although a small number of spectra were found for several other integrin subunits, primarily β1, the α subunits did not overlap between the two cell lines. In RPMI-7951 cells, but not in MDA-MB-435S cells, MS analysis identified integrin α5, which is in line with our own results that these cells form FBs (Stojanović et al., 2025) and can be found in RAs isolates because FBs are less dependent on actin cytoskeleton.

Talin2 was found to be not needed for RA maintenance, which indicates that talin2 is not an activator of integrins in RA. Potential activators of integrin β5 in RA include the proteins Dab2, Numb, and ARH, which have been shown to interact directly with integrin β5 (Calderwood et al., 2003), and whose binding affinity for integrin β5 is several times higher than that of talin1 (Zuidema et al., 2022).

The key finding here is that KANK2 is a component of RAs in both melanoma cell lines. Not only was KANK2 enriched in samples of isolated RAs from both cell lines, those that form FBs and those that do not, but PLA analysis after integrin α5 knockdown (to deplete FBs) and CytoD treatment (to deplete FAs) also revealed close proximity between talin2 and KANK2. Therefore, given that talin2 and KANK2 were found in three different types of adhesions, this suggested that they may play a role in the interconversion. The fate of FAs and RAs has been suggested to be mainly connected via their shared use of αVβ5 integrins. Manipulations that inhibited one of these structures appeared to favour the other (Lock et al., 2018, Lukas et al., 2024) presumably by enlarging the available αVβ5 integrin pool (Lukas et al., 2024). Our results support these data. Namely, in MDA-MB-435S cell line talin2 from integrin αVβ5 FAs and KANK2 from CMSC have been implicated in maintaining actin-MT crosstalk. Therefore, knockdown of either of these proteins alters integrin αVβ5 FAs dynamics, resulting in enlarged αVβ5 FAs along with reduced cell migration (Loncaric et al., 2023). The enlarged αVβ5 FAs suggest increased levels of FA proteins. Therefore, it was not surprising that knockdown of either talin2 or KANK2 led to reduced abundance of RA components. In RPMI-7951 cells, which form three talin2 and KANK2 positive adhesions (FAs, FBs and RAs), talin2 knockdown also increased FA size (suggestive of increased levels of FA proteins) and reduced cell migration (Stojanović et al., 2025) and again, a reduction in RAs levels was observed. However, in the RPMI-7951 cell line, KANK2 knockdown did not affect integrin αVβ5 FAs. Therefore, the unaltered αVβ5 FAs, likely explains why the abundance of RA components after KANK2 knockdown remain unchanged.

In MDA-MB-435S cells (which lack FBs), talin2 knockdown did not affect KANK2 levels, although the characteristic RA proteins were reduced. In RPMI-7951 cells, the very low number of KANK2-specific spectra made it impossible to determine whether KANK2 levels change; however, a clear decrease in specific RA proteins was observed. Thus, within RAs, talin2 depletion does not reduce KANK2 levels. Still, it remains unclear whether these two proteins directly interact within RAs and what functional role they might play. Conversely, in both melanoma cell lines, KANK2 knockdown reduced talin2 levels, regardless of the overall effect on RA-specific proteins. This finding suggests the talin2 dependence of KANK2 levels. Moreover, it is in line with the previously mentioned observation that RA formation is independent of both talin2, already observed by (Lock et al., 2018), and apparently, KANK2 (our own data).

Until recently, it was not clear how αVβ5 integrin physically transfers between FAs and RAs. It has been shown that RAs arise more frequently at sites where the FA marker paxillin was progressively replaced by stonin1. In addition, the conversions in the opposite direction also occur by the recruitment of paxillin onto pre-existing β5 integrin scaffolds. These data indicate that two integrin αVβ5-based adhesions, having distinct molecular compositions, are intimately linked and dynamically interconvertible (Lukas et al., 2024). FAs and RAs are inversely regulated complexes and the balance between these types of adhesions is influenced by phosphatidylinositols (Lock et al., 2018). High levels of activated myosin light chain (p-MLC) also correlated with integrin αVβ5 localizing to FAs over RAs (Zuidema et al., 2022). Decreased membrane tension together with formation of contractile stress fibers, promotes the assembly of FAs, reduces RAs, and decreased cancer cell migration (Djakbarova et al., 2024). In MDA-MB-435S cells talin2 knockdown enhanced stress fiber formation and reduced cell migration. In addition, talin2 knockdown increased the size of FAs indicating preference of αVβ5 for FAs over RAs (Loncaric et al., 2023). Therefore, our results are consistent with this mechanism of inverse regulation of FA and RA. These conclusions are further supported by results in the RPMI-7951 cell line, in which only talin2 alters FAs and consequently alters RAs.

This comprehensive analysis of RAs in two melanoma cell lines expands on previous findings and highlights novel molecular components. The presence of talin2 in RAs is confirmed and KANK2 is identified as a novel RA-associated protein in both cell lines. While neither protein is essential for RA formation, their knockdown significantly alters RA abundance and composition, reflecting their broader roles in other adhesion types such as FAs and FBs. These findings emphasize the importance of adhesion crosstalk and the dynamic balance between different adhesion structures, which is influenced by integrin composition, cytoskeletal integrity, and membrane tension.

## Author contributions

Material preparation, data collection and analysis were performed by [AR], [ML], [NS], [MF], [MR], [DH], and [AA-R]. [AR], [ML] and [NS] contributed equally to the experimental work. The first draft of the manuscript was written by [NS] and [AA-R], corrections were made by [AR], [ML], [JDH] and [MJH]. All authors read and approved the final manuscript. [NS] and [AA-R] equally contributed to the study conception and design.

## Availability of data and materials

All relevant data could be obtained from the corresponding author.

