## Supplementary Table S1 for "Regulation of reticular adhesions by KANK2 and talin2 in two melanoma cell lines"

### Additional file 1

[1]Laboratory for Cell Biology and Signalling, Division of Molecular Biology, Ruđer Bošković Institute, Zagreb, Croatia; [2]Manchester Cell-Matrix Centre, Faculty of Biology, Medicine & Health, University of Manchester, Manchester, United Kingdom; [3]Laboratory for Computational Biology and Translational Medicine, Division of Electronics, Ruđer Bošković Institute, Zagreb, Croatia; [4]Department of Life Science, Manchester Metropolitan University, Manchester, United Kingdom

*equal contribution

**Table S1**

**Table S1.** List of used antibodies and dyes.

| **WESTERN BLOT** | | | | | |
|---|---|---|---|---|---|
| ***Primary antibodies*** | ***Ref. No.*** | ***Distributor*** | ***Monoclonal/polyclonal*** | ***Species*** | ***Dilution*** |
| Anti-Filamin B | ab97457 | Abcam, USA | Monoclonal | Rabbit | 1:1000 in 5% milk |
| Anti-human talin1 | MCA4770GA | Bio-Rad, USA | Monoclonal | Mouse | 1:1000 in 5% milk |
| Anti-a-Adaptin 1/2 (C-8) AP-2 | sc-17771 | Santa Cruz Biotechnology, USA | Polyclonal | Mouse | 1:1000 in 5% milk |
| Anti-Integrin β5 | D24A5 | Cell Signaling Technology, USA | Monoclonal | Mouse | 1:1000 in 5% milk |
| Anti-Numb (C29G11) | #2756 | Cell Signaling Technology, USA | Monoclonal | Rabbit | 1:1000 in 5% milk |
| Anti-LDH | sc33781 | Santa Cruz Biotechnology, USA | Polyclonal | Rabbit | 1:400 in 5% milk |
| Anti-human talin2 | MCA4771GA | Bio-Rad, USA | Monoclonal | Mouse | 1:1000 in 5% milk |
| Anti-KANK2 | HPA015643 | Sigma-Aldrich, USA | Polyclonal | Rabbit | 1:1000 in 5% milk |
| ***Secondary antibodies*** | ***Ref. No.*** | ***Distributor*** | ***Monoclonal/ polyclonal*** | ***Species*** | ***Dilution*** |
| Goat anti-rabbit IgG (H+L) | 31466 | Invitrogen, USA | Polyclonal | Goat | 1:5000 in 5% milk |
| Goat anti-mouse IgG (H+L) | G21040 | Invitrogen, USA | Polyclonal | Goat | 1:10 000 in 5% milk |
| **IMMUNOFLUORESCENCE** | | | | | |
| ***Primary antibodies*** | ***Ref. No.*** | ***Distributor*** | ***Monoclonal/ polyclonal*** | ***Species*** | ***Dilution*** |
| Anti-Integrin β5 | D24A5 | Cell Signaling Technology, USA | Monoclonal | Mouse | 1:600 in 5% BSA |
| Anti-vinculin | ab129002 | Abcam, USA | Monoclonal | Rabbit | 1:100 in 5% BSA |
| Anti-human talin2 | MCA4771GA | Bio-Rad, USA | Monoclonal | Mouse | 1:100 in 5% BSA |
| Anti-KANK2 | HPA015643 | Sigma-Aldrich, USA | Polyclonal | Rabbit | 1:100 in 5% BSA |
| Anti-Integrin α5 | NBP2-50146 | Novus Biologicals, USA | Monoclonal | Mouse | 1:500 in 5% BSA |
| Anti-Numb (C29G11) | #2756 | Cell Signaling Technology, USA | Monoclonal | Rabbit | 1:600 in 5% BSA |
| Recombinant Alexa Fluor® 647 Anti-Vinculin | ab196579 | Abcam, UK | Monoclonal | Rabbit | 1:200 in 5% BSA |
| ***Secondary antibodies*** | ***Ref. No.*** | ***Distributor*** | ***Monoclonal/polyclonal*** | ***Species*** | ***Dilution*** |
| Anti-Mouse IgG Alexa Fluor 546 | A-11030 | Invitrogen, USA | Polyclonal | Goat | 1:1000 in 5% BSA |
| Anti-Mouse IgG Alexa Fluor 488 | #4408 | Cell Signaling Technology, USA | | Goat | 1:1000 in 5% BSA |
| Anti-Mouse IgG Alexa Fluor 405 | A-31553 | Invitrogen, USA | Polyclonal | Goat | 1:250 in 5% BSA |

| | | | | | |
|---|---|---|---|---|---|
| Anti-Rabbit IgG Alexa Flour, 555 | A-31572 | Invitrogen, USA | Polyclonal | Donkey | 1:1000 in 5% BSA |
| Anti-Rabbit IgG Alexa Fluor 647 | #4414 | Cell Signaling Technology, USA | Polyclonal | Goat | 1:1000 in 5% BSA |
| Anti-Mouse IgG1 Alexa Fluor 555 | A-21127 | Invitrogen, USA | Polyclonal | Goat | 1:1000 in 5% BSA |
| Anti-Mouse $IgG2_b$ Alexa Fluor 488 | A-21141 | Invitrogen, USA | Polyclonal | Goat | 1:1000 in 5% BSA |
| ***Dyes*** | ***Ref. No.*** | ***Distributor*** | | | ***Dilution*** |
| Phalloidin, Alexa Fluor 488 | P5282 | Sigma Aldrich, USA | | | 1:100 in 5% BSA |
| **PLA** | | | | | |
| ***Primary antibodies*** | ***Ref. No.*** | ***Distributor*** | ***Monoclonal/polyclonal*** | ***Species*** | ***Dilution*** |
| Anti-human talin2 | MCA4771GA | Bio-Rad, USA | Monoclonal | Mouse | 1:4000 in diluent* |
| Anti-KANK2 | HPA015643 | Sigma-Aldrich, USA | Polyclonal | Rabbit | 1:4000 in diluent* |
| ***Secondary antibodies*** | ***Ref. No.*** | ***Distributor*** | ***Monoclonal/ polyclonal*** | ***Species*** | ***Dilution*** |
| Navenibody M1 (40X) | NB.1.100.06 | Navinci Diagnostics AB, Sweden | | Mouse | 1:40 in diluent* |
| Navenibody R2 (40X) | NB.1.100.07 | Navinci Diagnostics AB, Sweden | | Rabbit | 1:40 in diluent* |

** part of NaveniFlexTM Cell MR kit*
