## Supplementary Figures S1-S6 for "Regulation of reticular adhesions by KANK2 and talin2 in two melanoma cell lines"

**Additional file 2**

[1]Laboratory for Cell Biology and Signalling, Division of Molecular Biology, Ruđer Bošković Institute, Zagreb, Croatia; [2]Manchester Cell-Matrix Centre, Faculty of Biology, Medicine & Health, University of Manchester, Manchester, United Kingdom; [3]Laboratory for Computational Biology and Translational Medicine, Division of Electronics, Ruđer Bošković Institute, Zagreb, Croatia; [4]Department of Life Science, Manchester Metropolitan University, Manchester, United Kingdom

*equal contribution

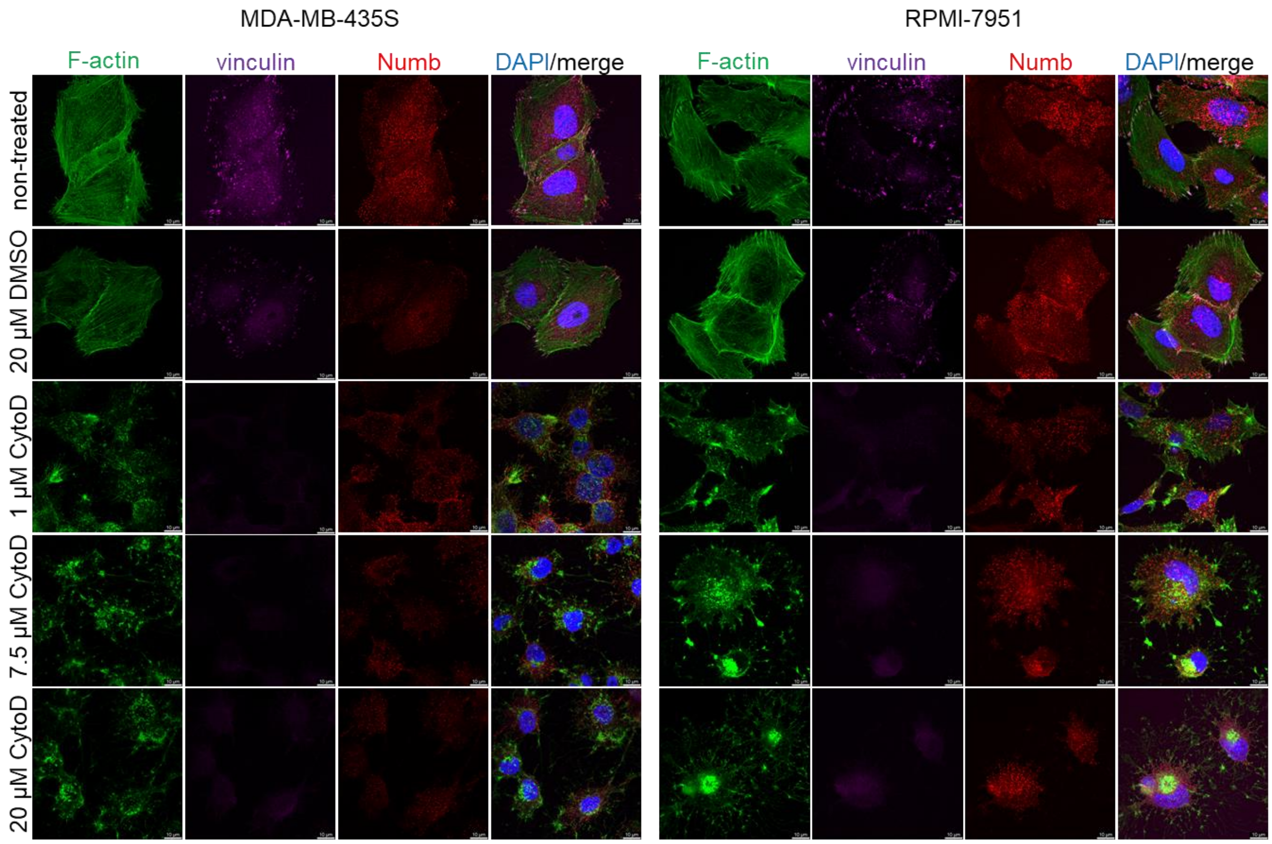


**Suppl. Figure S1** Optimization of CytoD concentration for enriching RAs in MDA-MB-435S and RPMI-7951 cells. Forty-eight hours upon seeding on coverslips, cells were treated with different concentrations of CytoD, fixed with PFA, permeabilised and stained with anti-vinculin and anti-Numb, followed by Alexa-Fluor 647-conjugated antibody (magenta) or Alexa-Fluor 546-conjugated antibody (red), respectively. F-actin staining (shown in green) was performed and IRM images were taken. Controls included non-treated cells and cells treated with a cytoD solvent DMSO. Analysis was performed using TCS SP8 Leica. Scale bar = 10 µm.

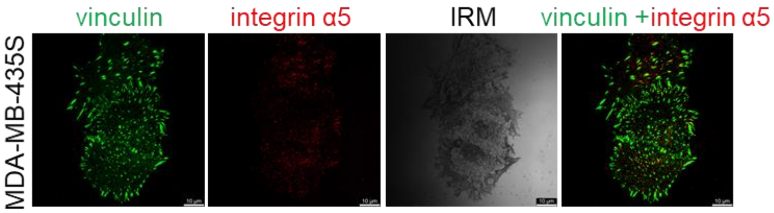


**Suppl. Figure S2** MDA-MB-435S cells contain very little to none α5-positive adhesions. Forty-eight hours upon seeding on coverslips, cells were fixed with PFA, permeabilised and stained with anti-vinculin and anti-integrin α5 antibody followed by Alexa-Fluor 488-conjugated antibody (green) or Alexa-Fluor 546-conjugated antibody (red), respectively. IRM images were taken. Analysis was performed using TCS SP8 Leica. Scale bar = 10 μm.

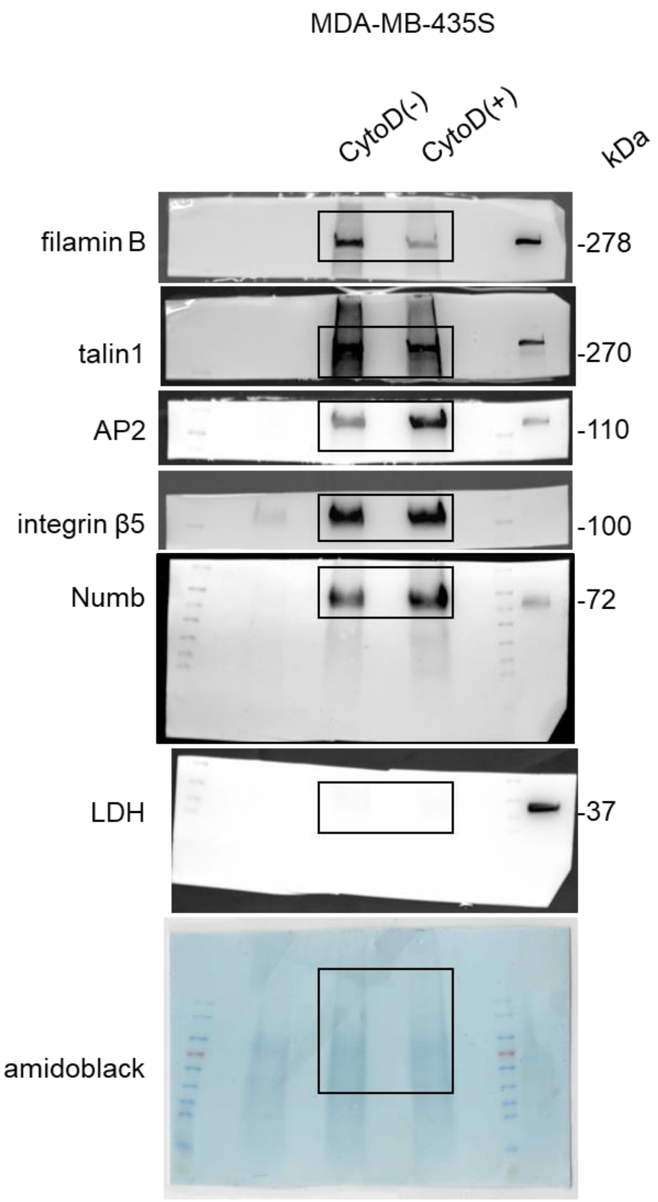


**Suppl. Figure S3** Full images of the blots in Figure 2C. Images were obtained using Uvitec Alliance Q9 mini, which directly scanned membranes developed with ECL reagent.

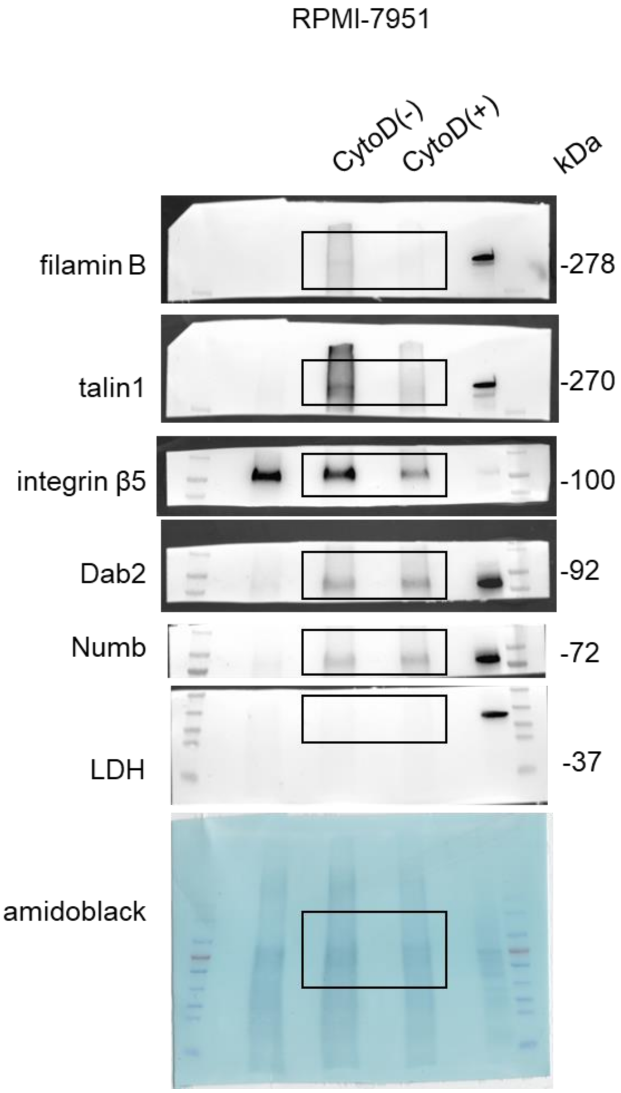


**Suppl. Figure S4** Full images of the blots in Figure 2D. Images were obtained using Uvitec Alliance Q9 mini, which directly scanned membranes developed with ECL reagent.

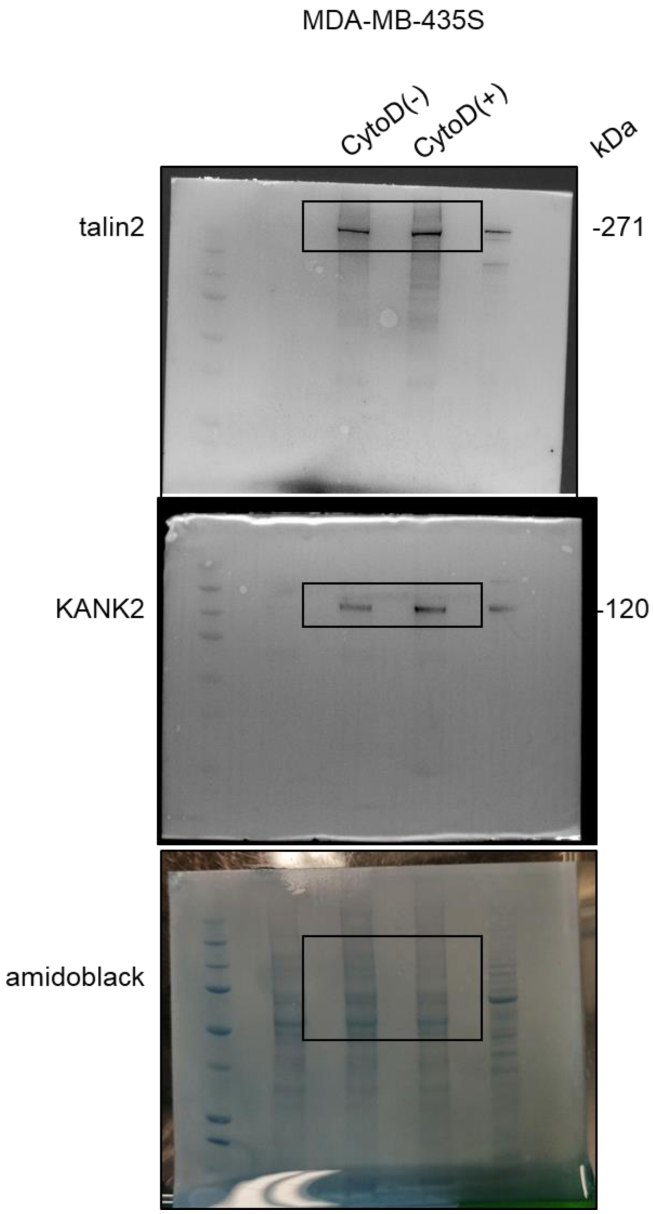


**Suppl. Figure S5** Full images of the blots in Figure 3B. Images were obtained using Uvitec Alliance Q9 mini, which directly scanned membranes developed with ECL reagent.

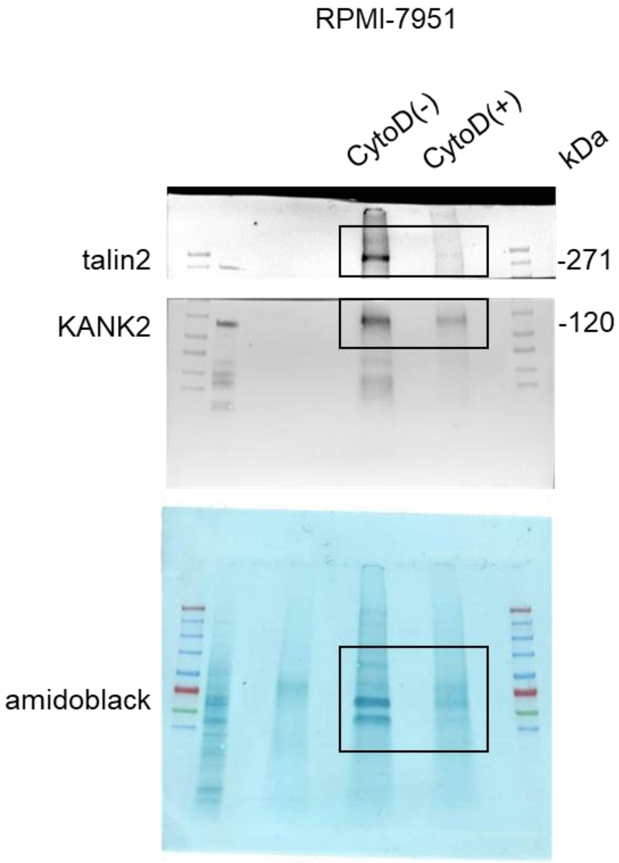


**Suppl. Figure S6** Full images of the blots in Figure 3C. Images were obtained using Uvitec Alliance Q9 mini, which directly scanned membranes developed with ECL reagent.
